# Muscle Contraction is Essential for Tendon Healing and Muscle Function Recovery after Achilles Tendon Rupture and Surgical Repair

**DOI:** 10.1101/2024.05.01.592124

**Authors:** Moe Yoneno, Yuki Minegishi, Haruna Takahashi, Kei Takahata, Himari Miyamoto, Yuna Usami, Takanori Kokubun

## Abstract

Incomplete tendon healing and postponed muscle weakness after Achilles tendon rupture and surgical repair lead to poor performance in patient activities. Although the effectiveness of postoperative early functional rehabilitation has been proven, the priority and each effect of specific methods in early rehabilitation remain unclear. We hypothesized early muscle contraction exercises without joint motion would promote tendon healing and prevent calf muscle atrophy; in contrast, early static stretching after surgical repair would not contribute to tendon healing and induce calf muscle atrophy. C57Bl/6 mice underwent Achilles tendon rupture and suture repair, followed by different methods of post-surgery interventions: a non-exercise group, a Static stretching group, and an Electrical muscle stimulation group. 3 and 5 weeks after surgery, we assessed ex vivo tendon mechanical properties, collagen fiber alignment, and histological muscle properties. Electrical Muscle Stimulation restored the recovery of tendon mechanical properties and muscle strength more quickly than Static stretching. Static stretching had no additional effect on them compared to the non-exercise. Our results suggested that calf muscle contraction was essential as a post-surgery early functional rehabilitation to load tensile forces on tendons and improve Achilles tendon healing. Additionally, early muscle contractions naturally promote restoring muscle function after the rupture, but further research is needed to optimize muscle contraction protocols.

**Statement of Clinical Significance:** This study shows the importance of selecting appropriate exercise modalities to resolve imperfections in tendon healing and muscle recovery. The establishment of proper rehabilitation is expected to improve post-surgery outcomes.

**Study Design:** A controlled laboratory study.

## Introduction

Achilles tendon rupture (ATR) is one of the most common injuries and frequently occurs in sports activities.^1^ After ATR, the dysfunction of ankle plantar flexion and low-performance level remain semi-permanent.^2,3,4,5^ Surgical treatment allows early functional rehabilitation (EFR), and its resume may lead to early functional recovery.^6,7^ Moreover, it is essential to reacquire tendon tension and begin exercise as early as possible after surgery to minimize muscle atrophy. However, the current report showed no difference in the re-tear rate or functional prognosis between surgical and conservative treatment.^6^ Therefore, establishing effective EFR protocols based on scientific evidence improves post-surgery outcomes. Recent reports mentioned the changes in Achilles tendon (AT) properties, such as length and stiffness, in patients after surgical repair.^8–11^ Of course, calf muscle atrophy also remains long-term.^12^ Incomplete tendon healing and calf muscle weakness lead to ankle dysfunction; thus, many clinicians and researchers have focused on effective post-surgical rehabilitation protocol to avoid these risks.

Biologically, mechanical loading contributes to promoting tendon extracellular matrix synthesis.^13,14^ Its contributions to tendon healing also have been reported in several animal models. Completely removing load is detrimental to tendon maturation,^16^ therefore, mechanical loading restores the mechanical properties of tendons.^17,18^ Even though these reports explain the importance of EFR after surgery. Eliasson et al. clinically indicated that early weight bearing and ankle mobilization didn’t improve the mechanical properties of healing tendons,^19^ suggesting appropriate EFR is still unclear.

In the tendon healing process, there is a contradiction between the biological knowledge and clinical rehabilitation research outcomes. While most clinical studies focused on when to start weight bearing or ankle mobilization, ^12,20,21,22,24^ Schrepel et al.^23^ indicated that post-operative resistance exercise of plantar flexion improves the mechanical properties of healing tendons without its elongation in human patient subjects. The authors strongly suggested that higher tensile loads can enhance the healing quality of the tendon compared to weight bearing with cast immobilization or ankle mobilization. Further studies are needed to analyze the effect of the different mechanical loads on the tendon healing phase after reconstruction. The knowledge of these effects leads to improved ways of EFR and consistent definitions of “how to exercise” for tendon healing after surgery.

Additionally, some studies indicated that early functional mobilization didn’t improve calf muscle functional recovery after ATR-Surgical Repair. ^12,20,21,22^ During the immobilization phase after surgery, the EFR doesn’t contain the muscle strength exercise to prevent tendon elongation, which may lead to the dysfunction of calf muscle contraction. However, the difference in the amount of tensile loading didn’t relate to tendon lengthening.^19^ And, even if the tendons elongate at once, the shortening occurs in the later healing phase. ^24^ In contrast to some aspects that tensile loading to healing tendons leads to positive effects on tendon recovery, there is a lack of evidence to suggest whether avoidance of load on healing tendons by calf muscle contraction prevents tendon elongation. This is the factor that inhibits earlier muscle strength recovery.

Generally, after the cast is removed 2 weeks post-surgery, they receive stable orthosis with some heel wedges. At this time, their ankle has been kept in the plantarflex position. Following 4-6 weeks after changing the cast to orthosis, their ankle range of motion is extended toward dorsiflexion by gradually removing the heel wedges. In this process, both the calf muscle and AT are stretched. The effect of treatment focusing on such muscle-tendon dynamics has yet to be thoroughly investigated. This factor would be a key to solving the incomplete recovery of tendon and muscle function. Researching patients to solve these problems is difficult because of the large individual differences.

This study aimed to clarify the effects of individualized different types of exercise after ATR-Surgical Repair on the early phase of tendon healing and calf muscle function. We compared two intervention models in mice with dorsiflexion static stretching or plantar flexor isometric contraction. We hypothesized that muscle contraction would promote tendon healing and restore calf muscle function, whereas static stretching would not contribute to them.

## Methods

### Experimental Design

The University Animal Experiment Ethics Committee approved all experiments. We used 66 male C57BL/6 mice, aged 10 weeks, weighing 24 ± 3g. The left ATs of all the animals were surgically dissected and sutured. Firstly, 6 mice were used to evaluate AT length changes during static stretching. Then, the others were randomly divided into four groups with different post-surgical intervention methods: the non-exercise group (NE), the static stretching group (St), the electrical muscle stimulation group (EMS), and the Intact-operated group (Intact; the lateral hindlimb of NE) (Figure 1). Animals were bred in 2 per cage and given food and water *ad libitum*. The Room temperature was 23 ± 2 °C, with a 12-hour light-dark cycle. At 3 and 5 post-operative weeks, histological and biomechanical evaluations were performed.

**Figure 1.**
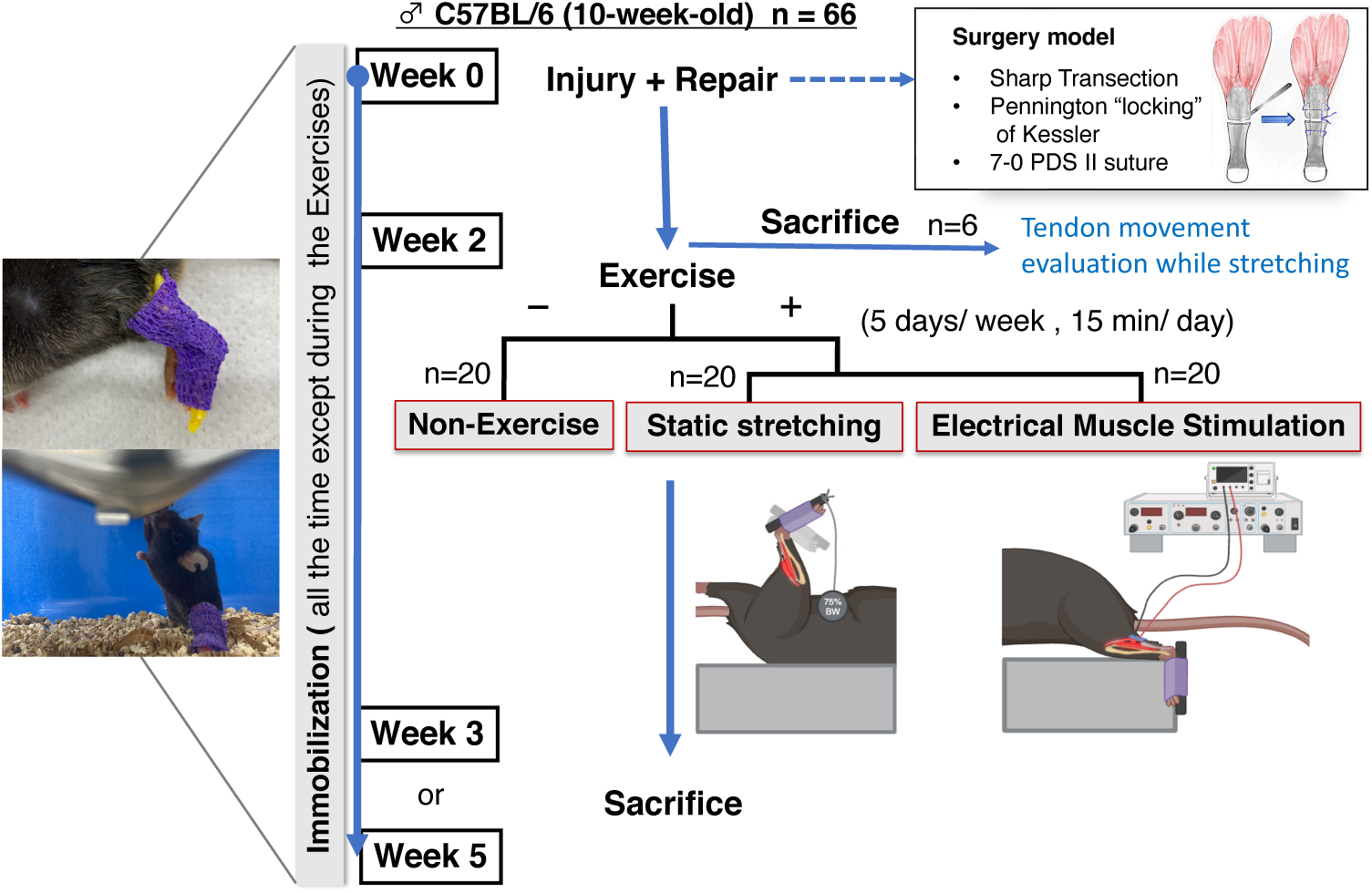
A flowchart showing the allocation of mice in this study. All mice were allowed weight-bearing with immobilization during their cage activities.

### Surgical procedure

Animals were given isoflurane inhalation anesthesia, followed by subcutaneous administration of a triad of anesthetics. As a pretreatment for surgery, lower limbs and feet were shaved and disinfected with 10% povidone-iodine. The surgical procedure was initiated after confirming the disappearance of escape reaction to stimulation of the toes with tweezers.

A longitudinal incision was made on the left lower limb with a No. 10 scalpel to expose the AT. After dissection of the middle part of the tendon with a No. 11 scalpel, the tendon was sutured by a 7-0 PDSⅡ suture (Johnson & Johnson). Their skin was closed with a 5-0 nylon suture (BEAR Medic). The contralateral hindlimb of all mice underwent only skin incision and closure.

After surgery, in all animals, their left ankle was fixed with the surgical intervention with external orthosis in the middle position. Then, they were awakened from anesthesia and allowed to move freely in the cage.

### Post-surgery Exercises

Exercise interventions were performed from 2 weeks after surgery and continued for 1-3 weeks, 5 days/ week. While exercising, all mice were anesthetized with 2% isoflurane inhalation.

#### NE group

Animals were not given additional exercise; they were only given continuous anesthesia in a plastic box.

#### St group

We used a custom-built passive ankle range of motion exercise device. 75 % of body weight was applied toward ankle dorsiflexion via a plantar plate with the foot fixed, and sustained stretching was performed continuously for 15 minutes.

#### EMS Group

We used a neuromuscular electrical stimulation device (Nihon Kohden, ), which set-up was based on a previous report by Ambrosio et al.^25^ The ankle joint was fixed at 90°, and the medial gastrocnemius muscle was stimulated via an electrode pad to perform isometric contraction. The stimulation duration was 15 minutes, with 6 sets of 10 stimulations per set and 5 minutes of rest between sets.

### AT length evaluation

6 animals were euthanized and exposed to the calf muscles and AT. Using the same system and setting animals in the same posture as in the St group intervention, we took photos at no load and 75% body weight load. From those photos, we evaluated AT length change using Image J Fiji.

### Tissue Sampling

We euthanized animals by carbon dioxide gas at the 1- and 3-week-exercising interventions (i.e., 3- and 5-weeks post-surgery). Then, gastrocnemius – AT – calcaneus complexities were harvested from 5 mice/ group. We dissected gastrocnemius at midpoint, and proximal parts were used to analyze muscle fiber morphometry. The muscles were immediately frozen in Hexane-Isopentane at -100℃. The distal portions of gastrocnemius were detached from the AT, and tendon–calcaneus specimens were used for biomechanical testing. We harvested the ankle joint from the other 5 mice/ group, including the healing tendon and the adjacent tissues, and analyzed the collagen fiber alignment of repair sights.

### Biomechanical Evaluation

We measured tendon thickness and width with a slide caliper before the testing. Cross-sectional area (CSA) was calculated as rectangles (thickness × width). ^26^ Tendons were then tested to evaluate the maximum force (N), stress (N/mm^2^), and stiffness (N/mm) with a mechanical testing machine (Univert, Cell Scale). Each specimen was preloaded at 0.05N and uniaxially tensioned (0.1mm/sec) until failure. Both ends of the specimens were wrapped with a soft microfiber wipe and cramped with the jigs. While testing, tendons were kept moisturized with phosphate-buffered saline.

### Tendon Collagen fiber Alignment

Each specimen was embedded in paraffin, sectioned sagittally at 5 µm-thick, and stained with picrosirius red to evaluate tendon collagen fiber alignment. After staining, we observed samples under a microscope (BZ-X700; KEYENCE) with a polarized light filter. Multiple bright-field and 90° circularly polarized images were taken at a magnification of ×20, and each image was merged to include the entire tendon. The region of interest (ROI; the healing area of the tendon) was selected from the bright-field image, and the rest of the image was masked to be excluded from the ROI. The ROI was superimposed on the polarized image, and 10 random 50 × 50 µm areas with no overlap were selected using Python 3.7. That average was calculated as the representative value in each sample. The more collagen is aligned parallel to the longitudinal axis, the higher the grayscale value, indicating more tendon maturity. ^27^

### Muscle fiber Morphometry

The frozen gastrocnemius muscles were sectioned at 10 µm-thick. To compare myofiber diameter and compositional changes of the myofiber types, multiple immunofluorescent staining of Myosin Heavy Chain (MHC) Type I, IIa, and IIb, and Laminin were performed based on the method of Ehmsen et al.^28^ The images were obtained with a BZ-X700 microscope. We used the image analysis software (Hybrid Cell Count; KEYENCE) to extract the inner diameter of the laminin-positive basement membrane as the myofiber perimeter. Then, we calculate the ferret diameter using Image J Fiji.^29^ The ferret diameter is the distance between parallel tangent lines of each myofiber perimeter. The mean value of the minimum ferret diameter of each fiber was used as the representative value of each sample for comparison between groups. The cross-sectional area of myofiber is proportional to the maximum muscle force, and the minimum ferret diameter is a geometric parameter that reliably predicts the cross-sectional area of myofibers. ^30^ The analysis was performed in the medial gastrocnemius muscle.

### Statistical Analysi**s**

Statistical analysis was performed using Python 3.7 (Python Software Foundation). The data were expressed as the mean ± SD. Our purpose was to compare specimens from 4 groups at each time point. All data were tested for normality by the Shapiro-Wilk test and equality of variance by the Bartlett test. Then, we analyzed by one-way ANOVA followed by the Tukey test for tendon grayscale score, transverse area, maximum rupture strength, stiffness, and muscle minimum ferret diameter.

Tendon stress was analyzed using a modified Welch test followed by the Games-Howell test. The statistical significance was set at *P* < .05.

## Results

### Tendon length evaluation

We compared the response of the tendon to passive stretching between the Achilles tendon suture and intact sides at 2-weeks after surgery. The results revealed load responses of the post-surgery tendons were similar to those of normal tendons. (Figure S1)

### Biomechanical Evaluation

Specimens from all mice failed at the middle substance close to the suture site. We showed the result of transverse tendon area (mm^2^), uniaxial peak force (N), stress (N/mm^2^), and stiffness (N/mm) testing at 3 and 5 week-post-surgery (Figure 2).

**Figure 2.**
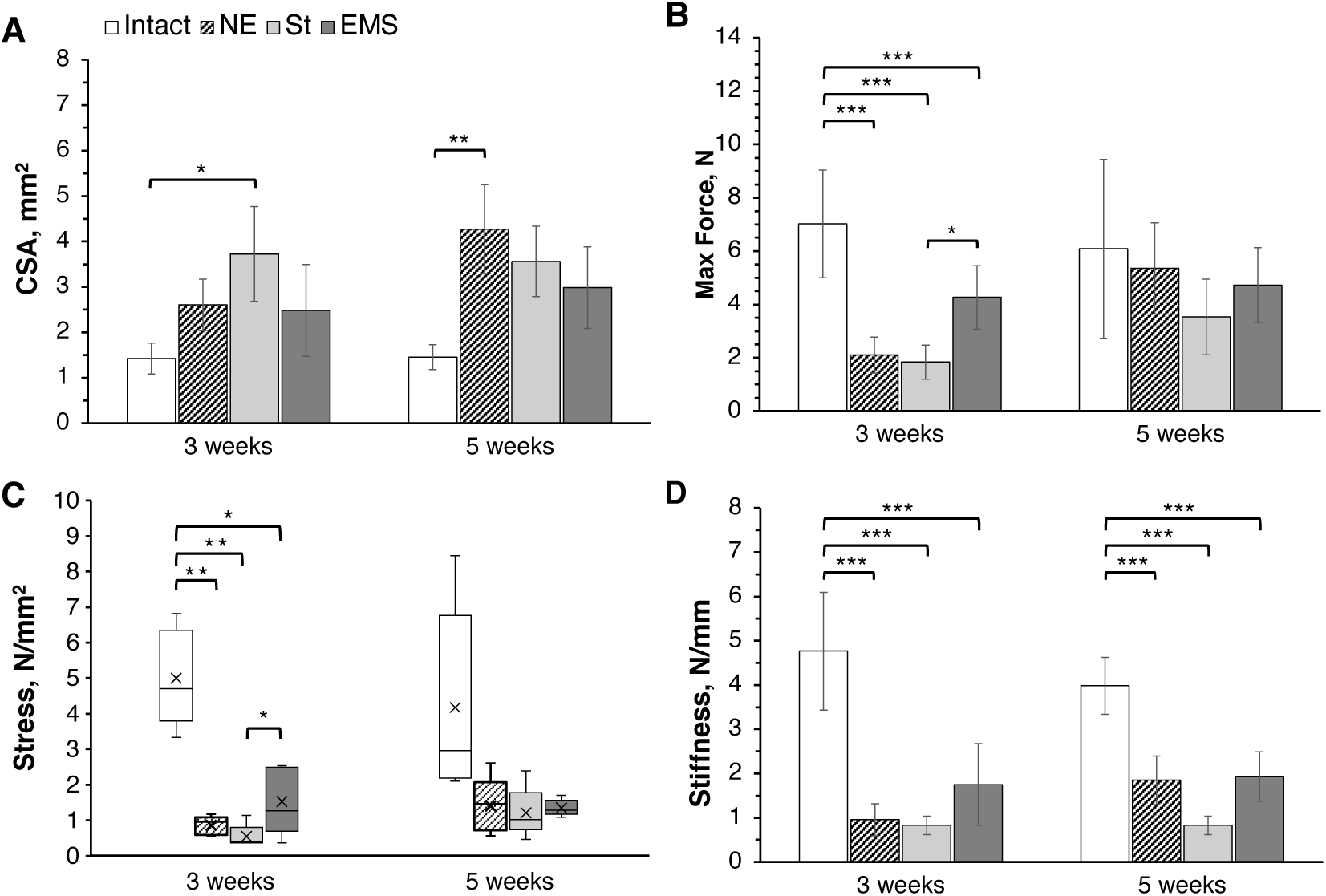
(A)The cross-sectional area of healing ATs at 3 and 5 weeks after surgery and their biomechanical properties: (B)maximum force, (C)stress, and (D)stiffness (\**P* < .05, \*\**P* <.01, \*\*\**P* < .001).

The CSA at 3 weeks in the St group was significantly higher than in the Intact. Those at 5 weeks in the NE group were significantly higher than in the Intact. The maximum force at three weeks was significantly lower in all the intervention groups than in the Intact. It was significantly higher in the EMS group than in the St group. At 5 weeks, there were no significant differences between the four groups. The stress at three weeks was significantly lower in all the intervention groups than in the Intact. It was significantly higher in the EMS group than in the St group. At 5 weeks, there were no significant differences between the four groups. The stiffness at three weeks was significantly lower in all the intervention groups than in the Intact. Still, there were no significant differences between the intervention groups. Those at 5 weeks were also lower in all the intervention groups than in the Intact. Still, there were no significant differences between the intervention groups.

### Tendon Collagen fiber Alignment

To evaluate the arrangement of collagen fibers in the healed tendons, we stained tendon sections with picrosirius red staining. In all intervention groups, collagen fibers were continuous and did not re-rupture (Figure 3-A). The grayscale score was calculated based on these images. It was significantly lower in all the intervention groups than in the Intact group. Still, there were no significant differences between intervention groups at three weeks and five weeks (Figure 3-B).

**Figure 3.**
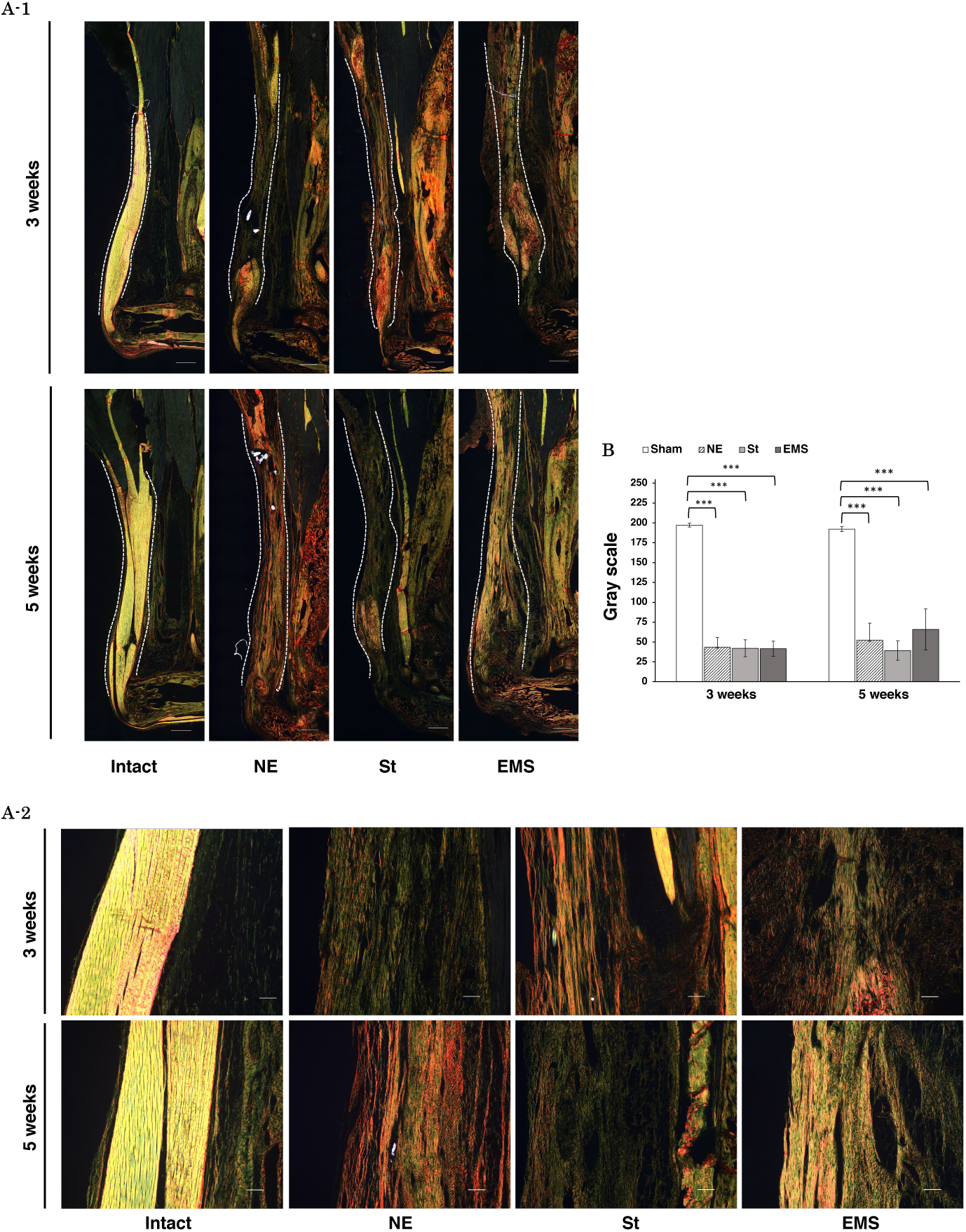
(A-1) Polarized view of the healing AT at 3 and 5 weeks after surgery stained with Picrosirius red. The collagen fibers were shown as orange, yellow, and green. ^31^ The images were taken with a 10× magnification lens and merged multiple (Scale bar = 500 μm). ATs were indicated between dotted lines. (A-2) The enlarged view of A-1 is surrounded by a pink square. (Scale bar = 100 μm) (B) The higher the grayscale score means, the more collagen fibers aligned parallel. (\*\*\**P* < .001)

### Muscle fiber Morphometry

We calculated the minimum Feret diameter per myofiber from immunofluorescence-stained images (Figure 4-A, B). They were significantly lower in all the intervention groups than in the Intact at 3 and 5 weeks. At three weeks, those were significantly higher in the EMS group than in the NE group. Still, there were no significant differences between the NE group and the St group and the St group and the EMS group. On the other hand, at 5 weeks, those in the EMS group were significantly lower than in both the NE group and the St group, while there were no significant differences between the St group and the NE group. In all the intervention groups, the minimum Feret diameter showed remarkable atrophy in Type Ⅱa and Ⅱx fibers (Figure 4-C), and the ratio of the TypeⅠfibers was decreased (Figure 4-D).

**Figure 4.**
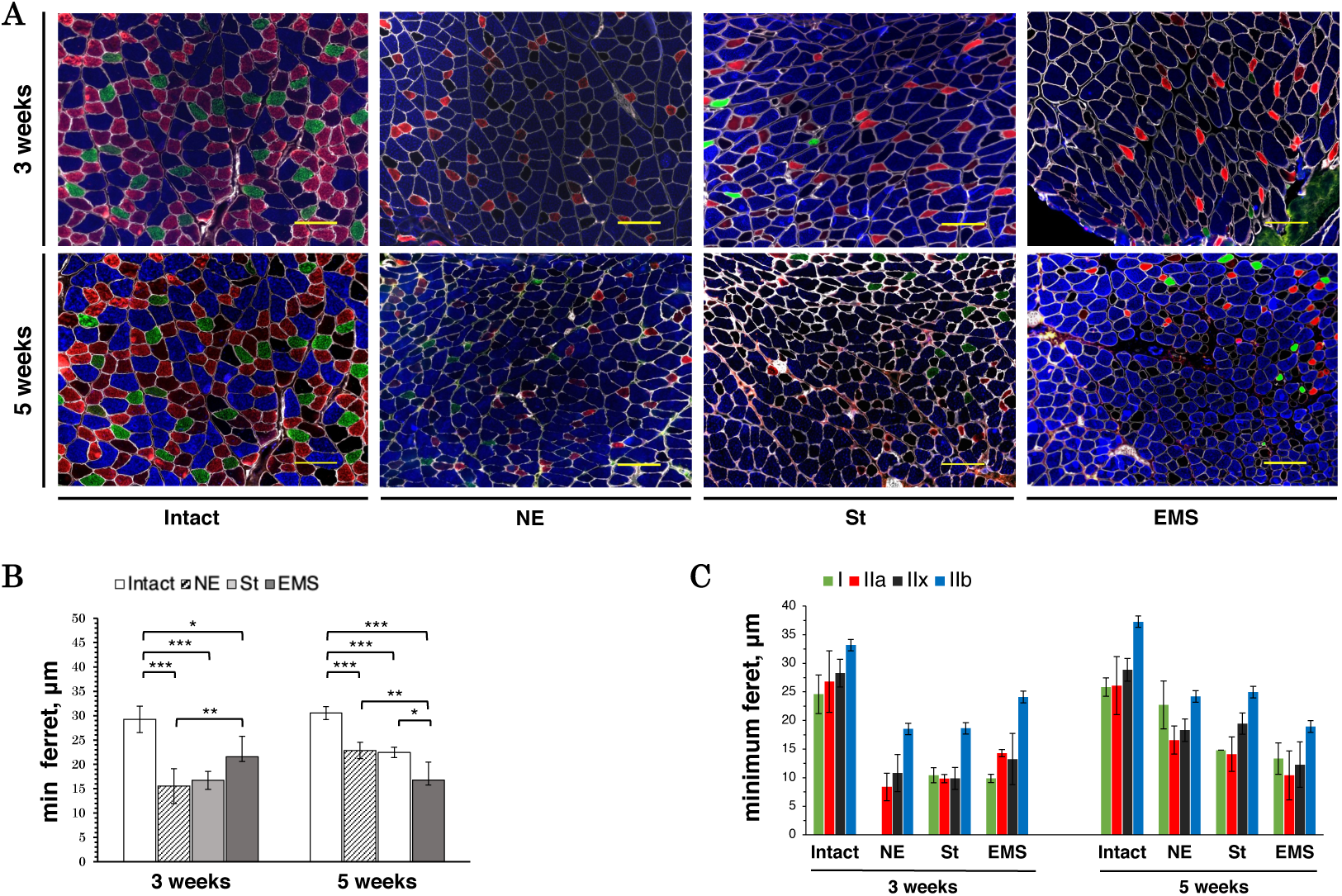
(A) Immunofluorescence (IF) images of medial gastrocnemius muscles at 3 and 5 weeks after surgery: MHC type Ⅰ (green), type Ⅱa (red), type Ⅱx (black), type Ⅱb (blue), and Laminin (white). The images were taken with a 20× magnification lens (Scale bar = 100 μm). (B) The minimum ferret diameters in all muscle fibers. (C) The minimum Feret diameter (mean ± SD) was calculated for each MHC type. Statistical analysis was not performed. (\**P* < .05, \*\**P* <.01, \*\*\**P* < .001).

## Discussion

This study aimed to assess the effectiveness of early mobilization and muscle contraction with AT healing and calf muscle function recovery after ATR-Surgical repair. We compared functional healing for three different groups: sedentary (NE), static stretch (St), and muscle strengthening exercise (EMS). The tendon mechanical properties in maximum force and stress showed higher recovery in the EMS group than the St group in earlier post-surgery phases. The result of healing tendon maturities showed no significant differences in grayscale between the three experimental groups at two-time points after surgery. Regarding muscle functions, the minimum Feret diameter of the gastrocnemius muscle fibers was significantly higher in the EMS group than the NE group at three weeks. However, at five weeks, the EMS group indicated a significantly smaller fiber size than the other two experimental groups. These results supported the importance of muscle contraction on AT healing and restoring muscle function in the early phase after surgical repair.

Ruptured tendons show diffuse enlargement of the tendon sheath and scar, including adhesion with surrounding tissues during the healing process.^32,33^ Scar-healed tendons have impaired mechanical properties and gliding function compared to natural tendons. While the increased tendon CSA compensates for mechanical strength,^15^ excessive scar formation leads to tendon dysfunction.^16^ Compared with the Intact, while the tendon CSA in the NE group at 5 weeks and the St group at three weeks were significantly increased, the EMS group had no significant difference. In addition, the EMS group showed better mechanical properties than the NE and St groups in terms of maximum failure force and stress. In the histological evaluation of the healing tendon, there was no significant difference in collagen maturation between the intervention groups at 3 and 5 weeks. However, the EMS group tended to have better collagen fiber orientation and maturation in the mid-substance of healing region at 5 weeks than the other two groups. In the general tendon healing process, fibroblasts were increased for several weeks after a few days of inflammation. Collagen type III is mainly synthesized in the healing region in the early healing phase. Then fibroblasts form a new extracellular matrix and replace thick, matured collagen type I.^34^ Previous studies showed this remodeling process continues over a few months to a year or more.^34–36^ This study obtained the histological data up to 5 weeks post-surgery. It was still in the process of tendon maturation. The critical finding here is that despite the higher mechanical loading to the tendon than other groups, EMS in the early phase did not inhibit the remodeling of its extracellular matrix but rather promoted qualitative recovery of its mechanical properties.

Excessive mechanical stress on sutured tendons may induce excessive inflammation and interfere with tendon healing; in contrast, the absence of mechanical stress also results in delayed remodeling. Recent studies reported that shortening the immobilization period and active exercise increase collagen fiber diameter and promote tendon maturation, and long-term ankle immobilization causes gaps, uneven thickness, and fragmentation of collagen fibers, resulting in disorder.^16,37–39^ In these studies, removing immobilization orthosis could increase mechanical loading on tendons due to a combination of factors, including the amount of loading, ankle range of motion, and muscle activity. Then, applying mechanical stress to the healing AT in some way contributes to tendon healing. In this study, postoperative ankle joint immobilization continued during their cage activities except for the exercise intervention time, following the immobilization period in clinical practice. Thus, mechanical loading on the musculotendinous generated mechanical loading during cage activity was restricted. This immobilization period is similar to the standard clinical protocol of 6 to 8 weeks. ^19,40^ The NE group did not have any additional exercise intervention. They were always immobilized in the mid-ankle position, which placed the least stress on the sutured tendon of the three intervention groups. In contrast, the St group was kept in the ankle dorsiflexion position with full extension of the knee joint during the intervention time, which means the entire calf muscle-AT complex was fully stretched. Still, the muscle was preferentially stretched during static stretching because the elastic modulus of the muscle was lower than that of the tendons.

On the other hand, in the EMS group, the muscles were stabilized in the mid-ankle position, and isometric contraction of the gastrocnemius muscle was performed. The muscle contractility forces directly generated the tensile stress on the sutured tendon. Based on comparing these two interventions, passive joint motion alone, as in the St group, was likely insufficient to transmit mechanical stress, promoting tendon healing. It can also be inferred that tendons were hardly stretched during the stretching intervention. In contrast, muscle contraction may have promoted higher-quality tendon healing, reducing adhesion to the surrounding tissues by sliding the tendon, preventing unnecessary scar formation, and promoting mechanical properties contributing to collagen fiber remodeling. These results suggest our hypothesis that muscle contraction exercises effectively promote tendon healing after ATR-Surgical Repair in the early phase after surgery.

Additionally, several reports mentioned that tendon lengthening occurs during the tendon healing process. Tendon lengthening induces slackness of muscle fiber. As a result, low plantar flexor function remains.^21,41,42^ So it is necessary to prevent tendon lengthening as much as possible. In the present study, the maximum tendon rupture strength and stiffness were not significantly different in the St and the EMS groups compared to the NE group. Since tendon stiffness represents the resistance of tendons to deformation, increased mechanical force by EMS on the healing tendon would not induce excessive tendon lengthening. Tendons in the EMS group were bearing to muscle contraction force, suggesting that subsequent muscle functional recovery may not affect tendon lengthening and be more efficient in restoring calf muscle function. Our results suggest that the tendon elongation mechanism in the AT repair healing process is not fitted by linear regression with the strength or amount of mechanical force. Further study is needed to assess the mechanism of exercise strategies for tendon healing during the early stages.

Regarding muscle recovery, we predicted that passive stretching would cause excessive muscle hyper lengthening and atrophy with decreased tensile strength. Contrary to our prediction, the NE and St groups showed no significant differences in minimum ferret diameters. On the other hand, the EMS group showed significantly greater muscle volume than the NE group but not different from the St group at 3 weeks. This result suggests that performing muscle contraction exercises in the early phase of the postoperative period can efficiently achieve both aims of tendon strength recovery and minimize muscle atrophy in preference to joint range-of-motion exercises. However, despite continued muscle strength training, muscle fiber diameters in the EMS group were significantly smaller than the NE and St groups at 6 weeks. The reason for this would be overload-induced microdamage of the calf muscle. The electrical stimulation parameters were the same as reported by Ambrosio et al..^25^ However, the number of intervention sets was increased to match the stretching time of the St group. The subject’s muscles had lost tension after AT rupture and atrophy at the beginning of the intervention due to post-surgery immobilization.^43^ Thus, continuous contractile exercise may have resulted in overload in the more vulnerable muscles. In addition, slow muscle fibers are preferentially mobilized by stimulation below 50 Hz.^44^ However, histological images and analysis of area by myofiber types showed a consistent decrease in Type I fibers and atrophy of Type II-a and II-x fibers in all the intervention groups. These results mean that the calf muscle shifted, increasing the slow muscle fiber after tendon injury. Against our expectation, the EMS group did not prevent those muscle morphological changes. McLachlan et al.^45^ showed the atrophy of slow-twitch fibers occurs within 24 hours after tendon rupture. Electrical muscle stimulation began 2 weeks after surgery in this study. That may have mobilized fast-twitch muscle fibers to compensate for the reduced responsiveness of slow-twitch muscle fibers, resulting in more fatigue. These factors may have caused muscle atrophy in the long-term intervention.

This study has several limitations. Firstly, since this study obtained the histological data up to 5 weeks post-surgery, it might have been necessary to observe a longer time for the maturation of collagen fibers to be reflected in the histological images. However, it was necessary to evaluate the effect of exercises on tendon healing during the early remodeling stage because once they degenerate, it’s difficult to restore the tendon properties later. Secondly, the protocol of electrical muscle stimulation for the EMS group was determined based on the previous study. ^25^ Calf muscles lose tension rapidly after tendon rupture. Thus, the protocol was probably excessive for them. The other limitation is that we didn’t evaluate the tendon elongation during the healing process. The residual tensile strength of our suture gets 60 % in 2 weeks and 40% in 4 weeks. We started each exercise intervention two weeks after surgery. We could evaluate that the exercise interventions didn’t cause re-rupture or inhibit the recovery of their mechanical properties. On the other hand, we didn’t show the final tendon length and whether it was related to muscle contractile function at 5 weeks.

However, our supplemental data suggest that dorsiflexion stretching preferentially stretches muscles, not tendons, and early stretching does not provide a beneficial extension load for tendon healing. It is conventionally believed that an early stretching of calf muscle induces tendon lengthening, starting with ankle immobilization at the plantar flexion position and gradually moving to the intermediate position. Static stretching in this study mimics this clinical practice in physical therapy. When a joint range of motion increases during the early period of post-surgery immobilization, it has conventionally been unclear what is being stretched. In short, conventional post-surgery treatment strategies may result in tendon weakness, lengthening, and insufficient muscle recovery because of inhibited maturation with little or no load on the tendon during the 6-8 week-long process.

In conclusion, muscle-strengthening exercise using electrical stimulation had a hypertrophic muscle effect after one week of intervention, did not interfere with tendon remodeling, and resulted in qualitative tendon recovery earlier than stretching exercises—however, overload-induced muscle fatigue and secondary muscle atrophy. Our results suggest that muscle contraction exercises may be more effective in promoting tendon healing and muscle recovery than the passive expansion of dorsiflexion ROM. Future studies will help to establish more appropriate EFR by examining the loading capacity of muscle contraction exercises and assessing muscle and tendon function in more detail. These results will help the clinical practice of physical rehabilitation in post-surgery outcomes.

## Acknowledgments

This work was supported by the Sasakawa Scientific Research Grant from The Japan Science Society (2021-6037).

