## Supplementary material for "Muscle Contraction is Essential for Tendon Healing and Muscle Function Recovery after Achilles Tendon Rupture and Surgical Repair": Figure S1

**Title**

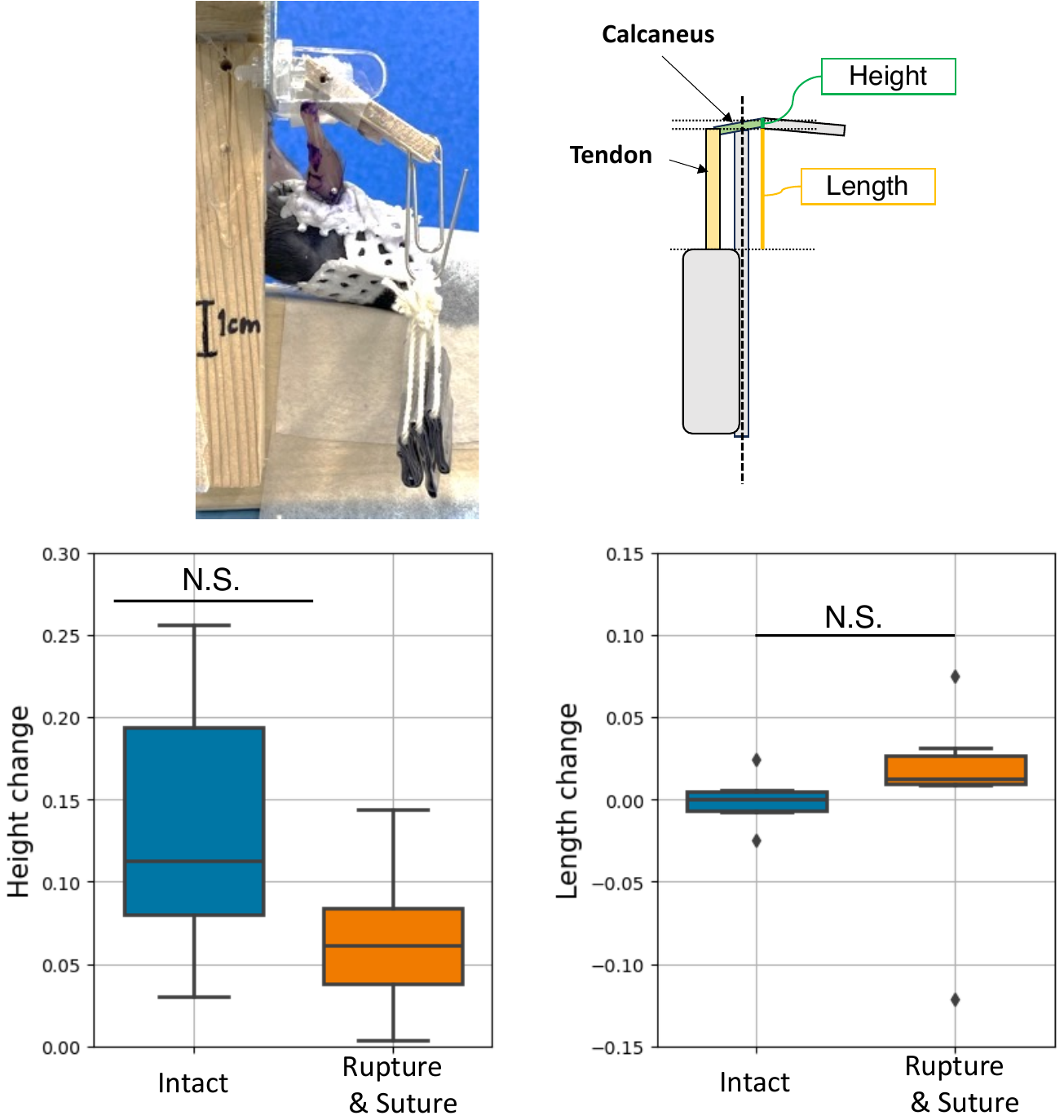

A

C

B

**Figure S1. Achilles tendon length evaluation while stretching**

(A) Pictures of measuring method. Calculated the tendon length and the calcaneus height difference before and after load. (B) Quantification of change of the calcaneus height when measuring tendon length (n = 6 mice). There were no significant differences between the Intact side and the Rupture & Suture side by unpaired, Mann-Whitney U-test (P < 0.05)*.*

(C) Quantification of change of tendon length and calcaneus height in mice 2 weeks after surgery. (n = 6 mice). There were also no significant differences between the Intact side and the Rupture & Suture side by unpaired, Mann-Whitney U-test (P < 0.05)*.*
